# Altered Relationship between Soluble TREM2 and Inflammatory Markers in Young Adults with Down Syndrome

**DOI:** 10.1101/776583

**Authors:** Grace E. Weber, Katherine Koenig, Maria Khrestian, Yvonne Shao, Elizabeth D. Tuason, Marie Gramm, Dennis Lal, James B. Leverenz, Lynn M. Bekris

## Abstract

Individuals with Down syndrome (DS) develop Alzheimer’s disease (AD) - related neuropathology, characterized by amyloid plaques with amyloid β (Aβ) and neurofibrillary tangles with tau accumulation more frequently and at an earlier age than their neurotypical counterparts. Peripheral inflammation and the innate immune response are elevated in DS. Triggering receptor expressed in myeloid cells 2 (TREM2) genetic variants are risk factors for AD and other neurodegenerative diseases. A soluble cleavage product of TREM2 (sTREM2) has been described as elevated in AD cerebrospinal fluid and positively correlates with Aβ and cognitive decline. There is relatively little information about TREM2 in DS. The objective of this study was to examine the relationship between sTREM2 and inflammatory markers in DS, prior to the development of dementia symptoms. Since TREM2 plays a role in the innate immune response and has been associated with dementia, the hypothesis of this exploratory study was that young adults with DS pre-dementia (n=15, mean age 29.5 years) would exhibit a different relationship between sTREM2 and inflammatory markers in plasma, compared to neurotypical, age-matched controls (n=16, mean age 29.6 years). Indeed, young adults with DS had significantly elevated plasma sTREM2 and inflammatory markers. In addition, in young adults with DS, sTREM2 correlated positively with 24 of the measured cytokines, while there were no significant correlations in the control group. Hierarchical clustering of sTREM2 and cytokine concentrations also differed between the group with DS and controls, supporting the hypothesis that its function is altered in people with DS pre-dementia. This exploratory study provides a basis for future studies investigating the relationship between TREM2 and the broader immune response pre-dementia.

## 1. Introduction

In the last half of the 20^th^ century, the life expectancy of individuals with Down syndrome (DS) has increased rapidly. The average lifespan is now 60 years ^1–3^, highlighting the importance of understanding age-related disease in this population ^4^. Most individuals with DS carry three full copies of chromosome 21. The amyloid precursor protein (*APP*) gene, located on chromosome 21, is processed into amyloid-β (Aβ). The overabundance of Aβ resulting from three copies of *APP* leads to Aβ plaque deposition in about 50% of people with DS over age 30 and subsequently increases their risk for dementia in middle age (around age 40) ^5,6^. The study of early adulthood in DS potentially provides a unique model for the early biology of AD in the general population.

Assessment of subtle cognitive changes is challenging in DS, partly because of the presence and variability of intellectual disability ^7,8^. Therefore, defining biomarkers of dementia progression are critical in this population of individuals. AD neuropathology, amyloid and tau, accumulates early in DS, and plasma Aβ and tau represent potential biomarker candidates ^9–11^. In DS, the variable severity of Aβ deposition within age groups and age of onset of dementia suggest that factors other than the presence of triplication of the *APP* gene contribute to the timing and severity of amyloid deposition and therefore the onset of AD ^5,12–14^.

Biomarkers of peripheral inflammation and the innate immune response are elevated in DS ^15–18^. Triggering receptor expressed in myeloid cells-2 (TREM2) is a pattern recognition receptor that can be activated by both pathogen- and danger- associated molecular patterns in the innate immune response ^19,20^. Loss-of-function genetic variants in *TREM2*, located on chromosome 6, are risk factors for AD and other neurodegenerative diseases ^21–23^. The soluble cleavage product of TREM2 (sTREM2) is elevated in AD cerebrospinal fluid and positively correlates with Aβ and cognitive decline ^24–32^. There is relatively little information about TREM2 in DS, though one group describes elevated sTREM2 in young adults and declining levels with aging ^33,34^.

The hypothesis of this investigation was that there is a relationship between soluble TREM2 (sTREM2) and inflammatory markers in DS, prior to the development of dementia symptoms. We found that young adults with Down syndrome displayed an altered immune profile compared to neurotypical controls, with increased levels of sTREM2 and inflammatory markers in plasma. In young adults with DS, sTREM2 correlated positively with 24 of the 38 measured cytokines, while there were no significant correlations in the control group. In addition, sTREM2 clustered with different cytokines in the two groups. The results of this exploratory study implicate an altered relationship between sTREM2 and inflammatory markers in young adults with DS pre-dementia.

## 2. Methods

### 2.1 Study Design

Fifteen young adults with DS (mean age 29.5 years, range 26-34 years; 5 females) and 16 neurotypical age-matched controls (mean age 29.6 years, range 25-36 years; 4 females) were enrolled in the study under a protocol approved by the Cleveland Clinic Institutional Review Board. All participants with DS had a previous medical diagnosis of DS ^35^, though results of chromosomal analysis were not available for all participants. Participants with DS provided verbal or written informed assent as appropriate, and their guardians provided written informed consent. Control participants provided written informed consent. Biospecimens were collected and processed through the Lou Ruvo Center for Brain Health Aging and Neurodegenerative Disease Biobank (LRCBH-Biobank). In order to minimize the chance of including participants that were showing subtle signs of dementia, each participant with DS underwent a comprehensive clinical assessment, including a psychiatric diagnostic interview with participant and caregiver.

### 2.2 Complete Blood Counts

Venipunctures were performed for the collection of whole blood, and plasma was isolated from lavender-top EDTA tubes. Complete blood counts (CBC) were performed on most participants (n=14 DS and n=15 controls) via microscopy by trained technicians blinded to sample group.

### 2.3 APOE4 genotyping

*APOE4* genotyping was performed from blood samples using the 7500 Real Time PCR System and TaqMan SNP Genotyping Assays (rs429358, rs7412) (Thermo Fisher) as previously described ^36^.

### 2.4 Plasma Biomarkers

Plasma soluble TREM2 (sTREM2) levels were measured using a Luminex 200 3.1 xPONENT System (EMD Millipore, Chicago, IL, USA) and a custom detection method designed to capture the soluble portion of TREM2 protein as previously described ^27^. Briefly, a capture antibody bound to MagPlex beads binds sTREM2 (R&D #MAB1828 human TREM2 antibody monoclonal mouse IgG_2B_ Clone #263602; Immunogen His19-Ser174). A biotinylated antibody with a SAPE conjugate was used for detection (R&D: #BAF1828; human TREM2 biotinylated antibody; antigen affinity-purified polyclonal goat IgG; Immunogen His19-Ser174). Inflammatory markers were measured with a human cytokine/chemokine panel utilizing Luminex 200 ×Map technology and the MILLIPLEX MAP ® multiplex kits (Luminex xMAP technology; EMD Millipore, Chicago, IL, USA: HNABTMAG-68K and HCYTMAG60PMX41BK, respectively) following the manufacturer’s instructions for analyte detection in human plasma. The inflammatory markers in the panel were: Epidermal Growth Factor (EGF), Fibroblast Growth Factor 2 (FGF-2), eotaxin, Transforming Growth Factor alpha (TGF-α), Granulocyte-colony stimulating factor (G-CSF), FMS-like tyrosine kinase 3 ligand (Flt-3L), Granulocyte-Macrophage Colony Stimulating Factor (GM-CSF), Fractalkine (also known as CX3CL1), interferon (IFN) alpha 2 (IFNα2), IFNγ, growth-regulated oncogene (GRO), interleukin (IL) 10, (IL-10), Monocyte chemotactic protein-3 (MCP-3), IL-12 40kDa (IL-12p40), Macrophage-derived chemokine (MDC), IL-12 70kDa (IL-12P70), IL-13, IL-15, soluble CD40-ligand (sCD40L), IL-17A, IL-1 receptor agonist (IL-1RA), IL-1α, IL-9, IL-1β, IL-2, IL-3, IL-4, IL-5, IL-6, IL-7, IL-8, interferon-gamma inducible protein 10 kDa (IP-10), Monocyte chemoattractant protein (MCP) 1 (MCP-1, also known as CCL2), MCP-3 (also known as CCL7), macrophage inflammatory protein (MIP) 1α (MIP-1α also known as CCL3), MIP-1β (also known as CCL4), tumor necrosis factor (TNF) alpha (TNFα), TNFβ), and Vascular Endothelial Growth Factor (VEGF).

### 2.5 Statistical Analyses

DS and control demographics were compared using unpaired t-test for age and Fisher’s exact test for *APOE*4 status and sex distribution. Plasma sTREM2 and inflammatory markers were measured in plasma on plates that also contained buffer alone background wells (containing zero pg/ml of each protein) and standard curve wells containing known concentrations of each protein measured. Concentrations of analytes were determined based on fluorescence of the standard curves for the respective proteins. Concentration values were log_10_ transformed for comparison. Values below fluorescence levels of the background wells were considered to be physiologically zero pg/ml, and set equal to 0.001pg/ml in order to be able to log transform the data. Individual plasma samples were run in duplicate on two different plates, and the overall mean of the 4 replicates was determined. Replicates with a high coefficient of variation (>30%) were analyzed to identify outliers, which were then removed from the analysis. CBC, biomarker, and inflammatory marker data were compared between groups using unpaired t-tests, and false discovery rates (FDR) were determined using the Original FDR method of Benjamini and Hochberg, with Q = 1%. Linear regression was used to determine correlations between analytes using SPSS version 22 (IBM Corp., Armonk, NY). Correlograms were made using the “corrplot” function in the corrplot package ^37^ and arranged according to hierarchical clustering using the Ward’s method in R version 3.6.1 (R Foundation for Statistical Computing, Vienna, Austria). Heat maps, including hierarchical clustering with the Ward’s method, were produced using the “heatmap.2” function in the gplots package ^38^ in R 3.6.1. Data were scaled using the “scale” function in R prior to input into the “corrplot” and “heatmap.2” functions. All other data analysis utilized GraphPad Prism version 8.1.1 (GraphPad Software, San Diego, CA).

## 1. Results

### 3.1 Cohort characteristics

Young adults with DS did not differ significantly from neurotypical controls in terms of age (mean 29.5 years vs. 29.6 years, p=0.89), distribution of sex (33.3% vs. 25.0%, p=0.70) or presence of the AD risk allele *APOE*4 (13.3% vs. 18.8%, p>0.99) (**Table 1**).

**Table 1.**
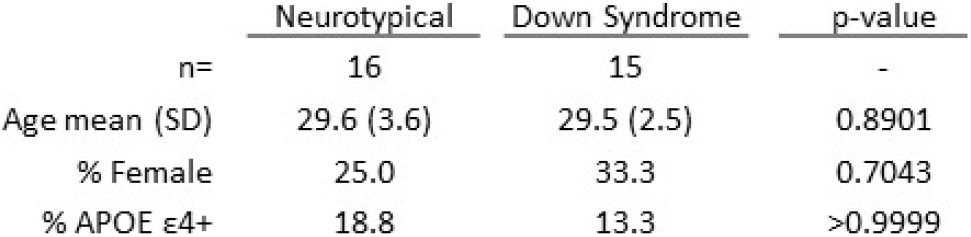
Population Description. Characteristics of the study populations were compared between neurotypica l controls (n=16) and ind ividuals with Down syndrome (n=15). There were no statistica lly s ignificant differences in age (unpa ired t-test), gender or APOE4 status (Fisher’s exact test) between groups.

|  | Neurotypical | Down Syndrome | p-value |
| --- | --- | --- | --- |
| n= | 16 | 15 | - |
| Age mean (SD) | 29.6 (3.6) | 29.5 (2.5) | 0.8901 |
| % Female | 25.0 | 33.3 | 0.7043 |
| % APOE ε4+ | 18.8 | 13.3 | >0.9999 |

### 3.2 Complete blood counts and inflammatory markers

Plasma sTREM2 was significantly elevated in DS compared to neurotypical controls (p=0.000966) **(Figure 1A).** Markers of inflammation, C-reactive protein (CRP) and erythrocyte sedimentation rate (ESR) were compared; CRP was elevated in DS (p=0.0169) and ESR trended toward being elevated in DS (p=0.05098) compared to controls **(Figure 1B-C).** Percentages and absolute values of cell types were compared between groups. Participants with DS had significantly higher percentages of basophils (p=0.000185) and trended towards higher percentages of neutrophils (p=0.087), compared to controls, while percentages of lymphocytes and monocytes did not differ between groups **(Figure 1D).** When comparing absolute cell counts, participants with DS had fewer lymphocytes (p=0.037) and more basophils (p=0.001) compared to controls **(Figure 1E).**

**Figure 1.**
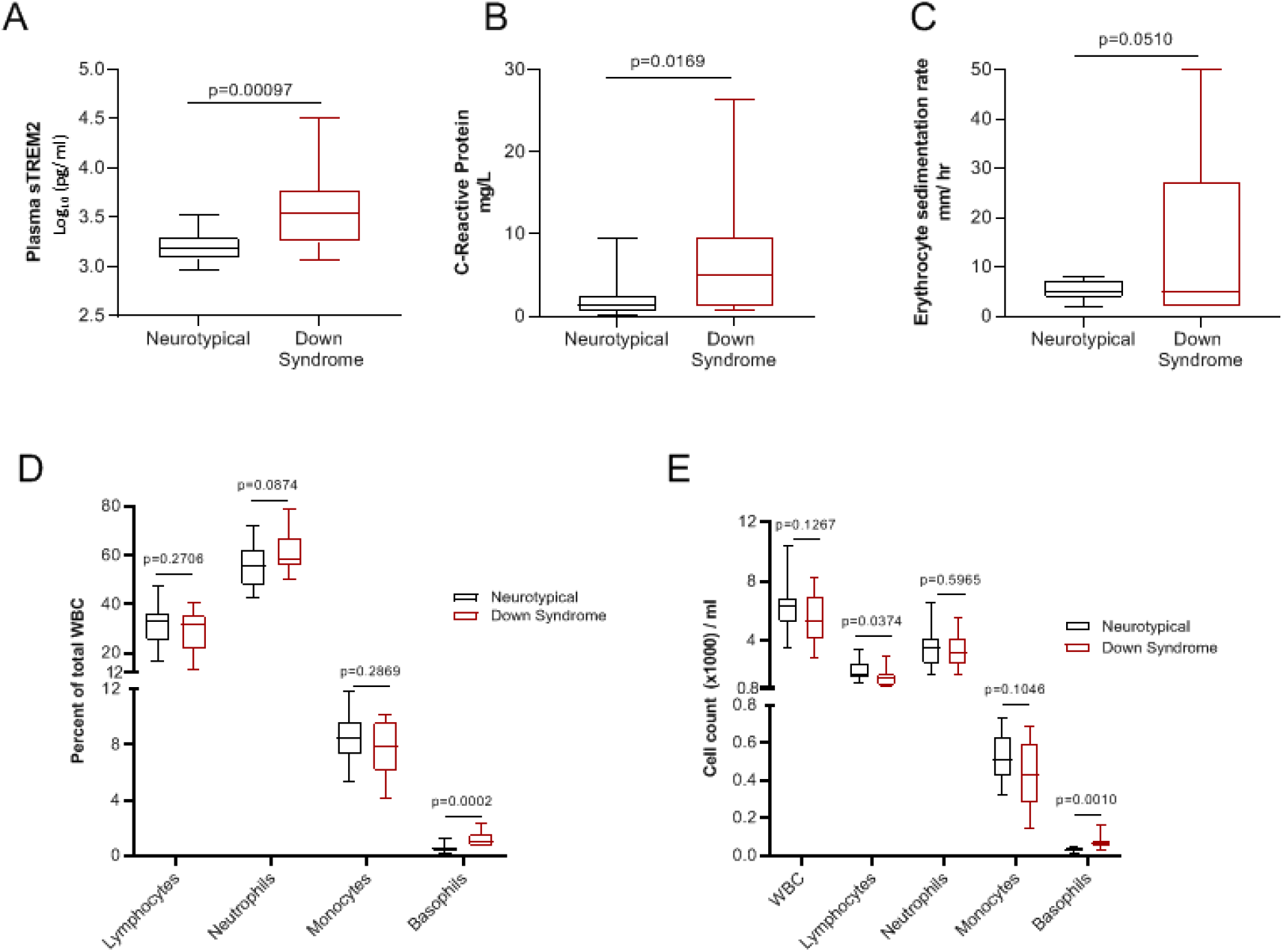
Immune markers and peripheral blood cell types. SignificantlyhigherplasmasTREM2 levelsin Down syndrome pre-dementia (n=15), compared to neurotypical (n=16) (p = 0.000966; 1% FDR significant) **(A)**. Inflammatory markers C-Reactive protein **(B)** and erythrocyte sedimentation rate **(C)** were elevated in OS compared to controls. White blood cell subset percentages **(D)** and cell counts **(E)** were compared between groups. Unpai red t-tests were performed between groups, p-vatuesare shown.

### 3.3 Levels of Plasma sTREM2 and Plasma Inflammatory Markers in Down Syndrome Pre-dementia

Given prior observations of elevated peripheral immune factors ^15,16,18,39^ and our own finding of elevated sTREM2 in DS, we addressed the question of whether other plasma inflammatory markers (cytokines, chemokines, and growth factors) were elevated in our DS cohort using an immune profiling panel. Inflammatory markers were grouped based on known function into the general categories “pro-inflammatory”, “immunoregulatory/ pleiotropic”, and “anti-inflammatory” ^40–49^. Values below assay background levels were determined to be physiologically zero pg/ml, and set = 0.001 pg/ml for the purpose of log transformation and analysis. Out of the 38 factors tested, 32 were significantly higher in DS compared to age-matched controls (**Figure 2A-B**); the exceptions were IL-4, FGF-2, MDC, IL-17A, GRO, and MCP-1. In order to determine if it was valid to set values below background equal to 0.001, we repeated the analysis after removing all values that had been below background detection levels (**Supplemental Figure 1A)**. We found that 30/38 cytokines were significantly higher in DS compared to controls; the exceptions were IL-13, G-CSF, IL-6, MDC, IL-17A, GRO, sCD40L, and MCP-1 (**Supplemental Figure 1AB)**.

**Figure 2.**
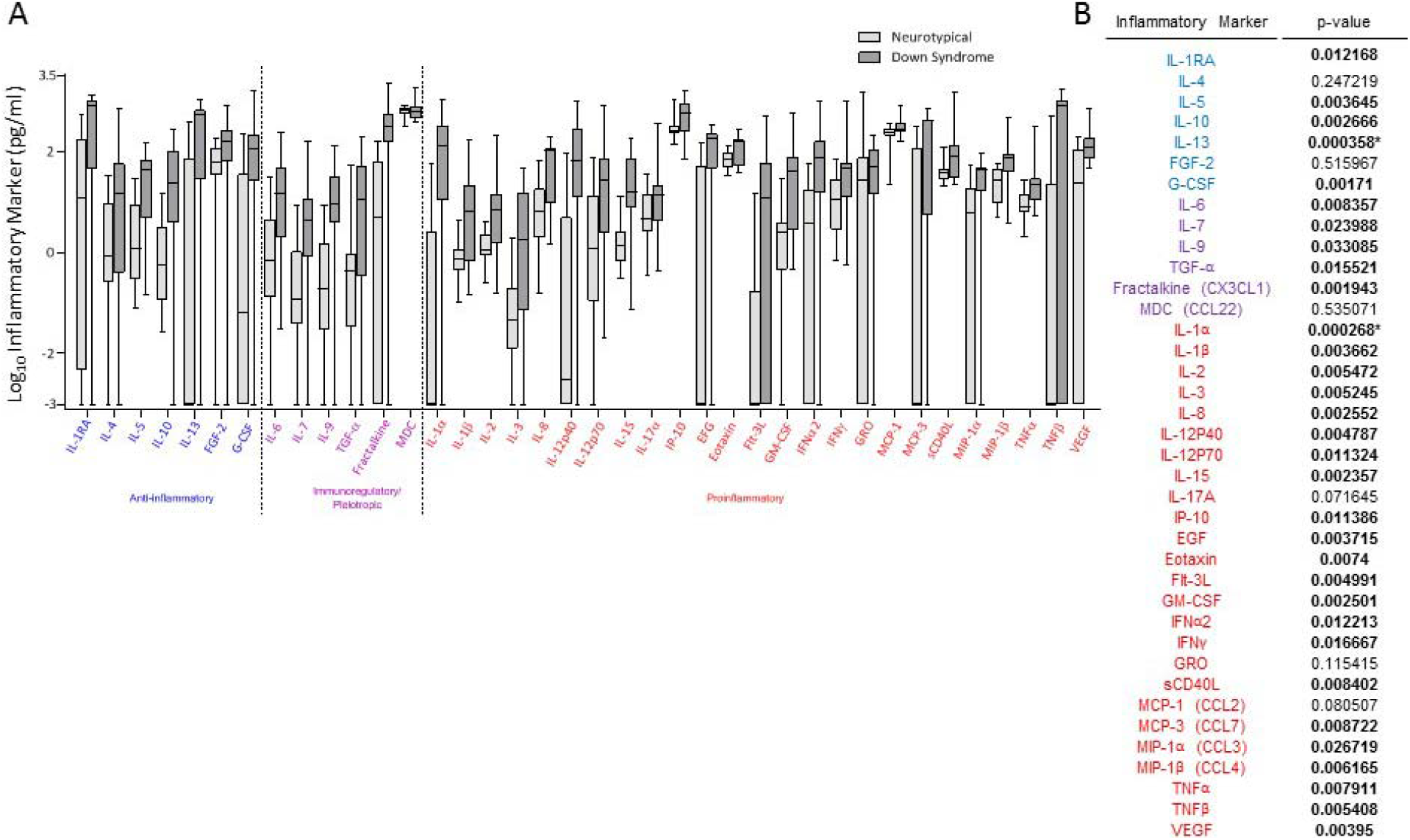
Plasma Inflammatory Markers in Down Syndrome Pre-dementia. Significantly higher plasma inflammatory markers in young adults with Down syndrome pre-dementia (n=15, dark grey bars), compared to neurotypical controls (n=16, light grey bars) for 32/38 of the inflammato ry marke rs on the panel **(A)**, significance as indicated by un paired t-test p-values (asterisk denotes 1% FDR significance) **(B)**. Inflammatory markers are separated by known functions into the general categories of anti-inflammatory (blue), immunoregulatory/pleiotropic (purple), and proinflammatory (red). Individual data points that had calculated concentrations below background levels for the assay were considered physiological Os and set equal to 0.001 prior to log transformation for analysis purposes.Lines indicate mean levels. Minimum and maximum levels are indicated.

### 3.4 The Relationship between Peripheral sTREM2 and Peripheral Inflammatory Markers

Pearson correlations were plotted and the correlation matrix was clustered based on the Ward’s method of least variance. The clustering showed distinctly different patterns in neurotypical controls compared to the DS group (**Figure 3A-B).** Importantly, sTREM2 showed statistically significant positive correlations only in the group with DS. In the neurotypical group, sTREM2 correlated negatively, though not significantly, with 17/38 cytokines, while in the DS group, sTREM2 did not correlate negatively with any cytokines. Instead, we observed significant positive correlations with sTREM2 and 24/38 measured cytokines in the DS group (MDC, sCD40L, TNFα, TGF α, IFNα2, IL-6, Flt-3L, GM-CSF, IL-1β, IL-12p70, IFNγ, MIP-1β, IL-17A, VEGF, IL-5, eotaxin, IL-8, IL-3, IL-4, IL-12p40, IL-9, IL-7, Il-10, and IL-15), and 0/38 measured cytokines in the control group (**Figure 3A-B**, Pearson correlation coefficients (r) and p-values are shown in **Supplemental Table 1**).

**Figure 3.**
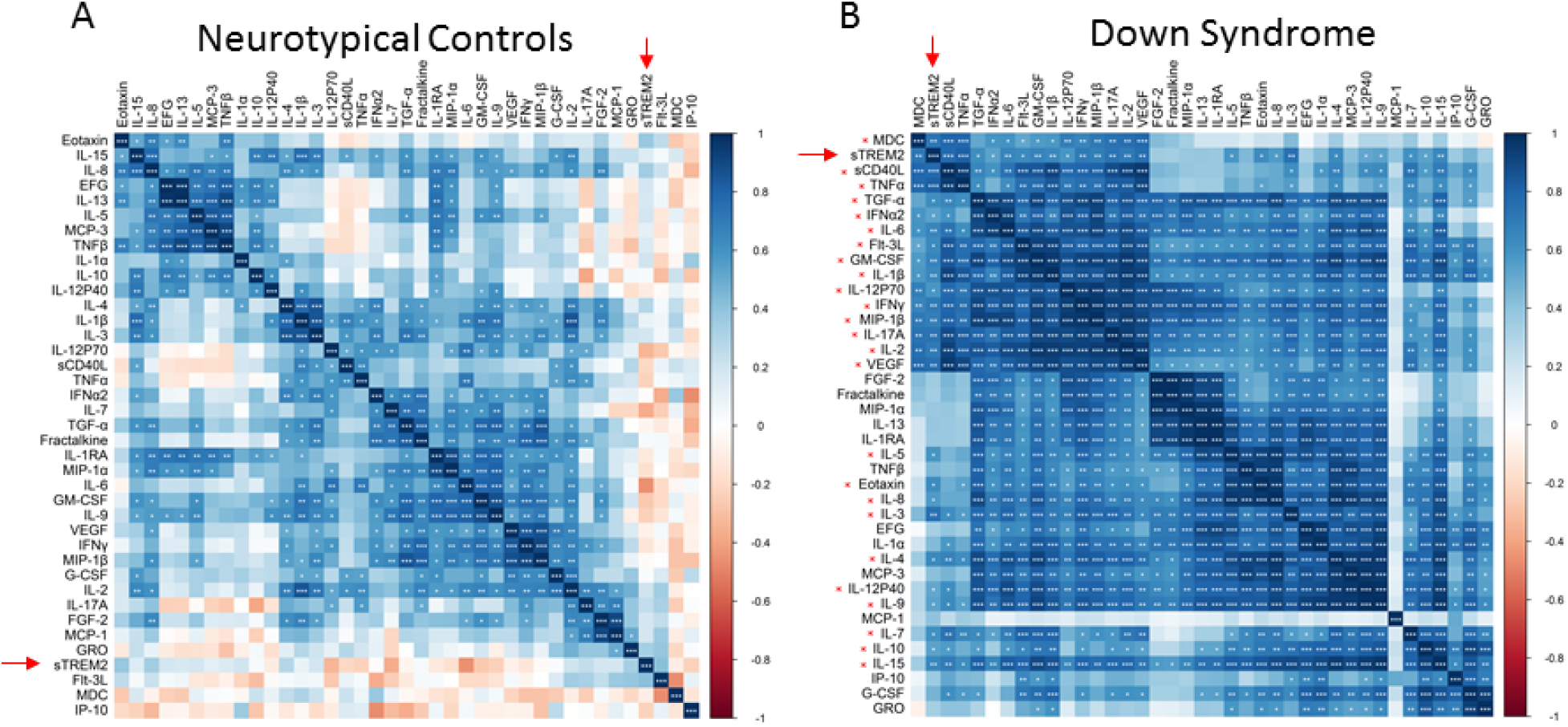
sTREM2 correlates positively with many inflammatory markers in DS. Pearson correlations of sTREM2 and inflammatory markers in plasma from neurotypical controls **(A)** versus young adults with Down syndrome **(B)**. Color gradient shows Pearson correlation coefficients (r), with dark blue = 1, indicating a perfect positive correlation, and red = −1, indicating a perfect negative correlation. The clustering patterns were determined by Ward’s method of least variance and differ between groups; negative correlations were seen only in the control group. sTREM2 is highlighted with a red arrow. Red asterisks next to inflammatory marker labels in the vertical orientation indicate significant correlation with sTREM2 of p<0.05. Asterisks within the boxes indicate significance of * p<0.05, ** p<0.01, *** p<0.001.

To further investigate this relationship, hierarchical clustering of the scaled log_10_ transformed immune factor concentration values was performed on each group. Lower sTREM2 levels in neurotypical controls (**Figure 1A**) clustered closest to GRO, Flt-3L, MDC, and IP-10, while higher sTREM2 in the DS group (**Figure 1A**) clustered with MDC, sCD40L, and TNF (**Figure 4A-B**).

**Figure 4.**
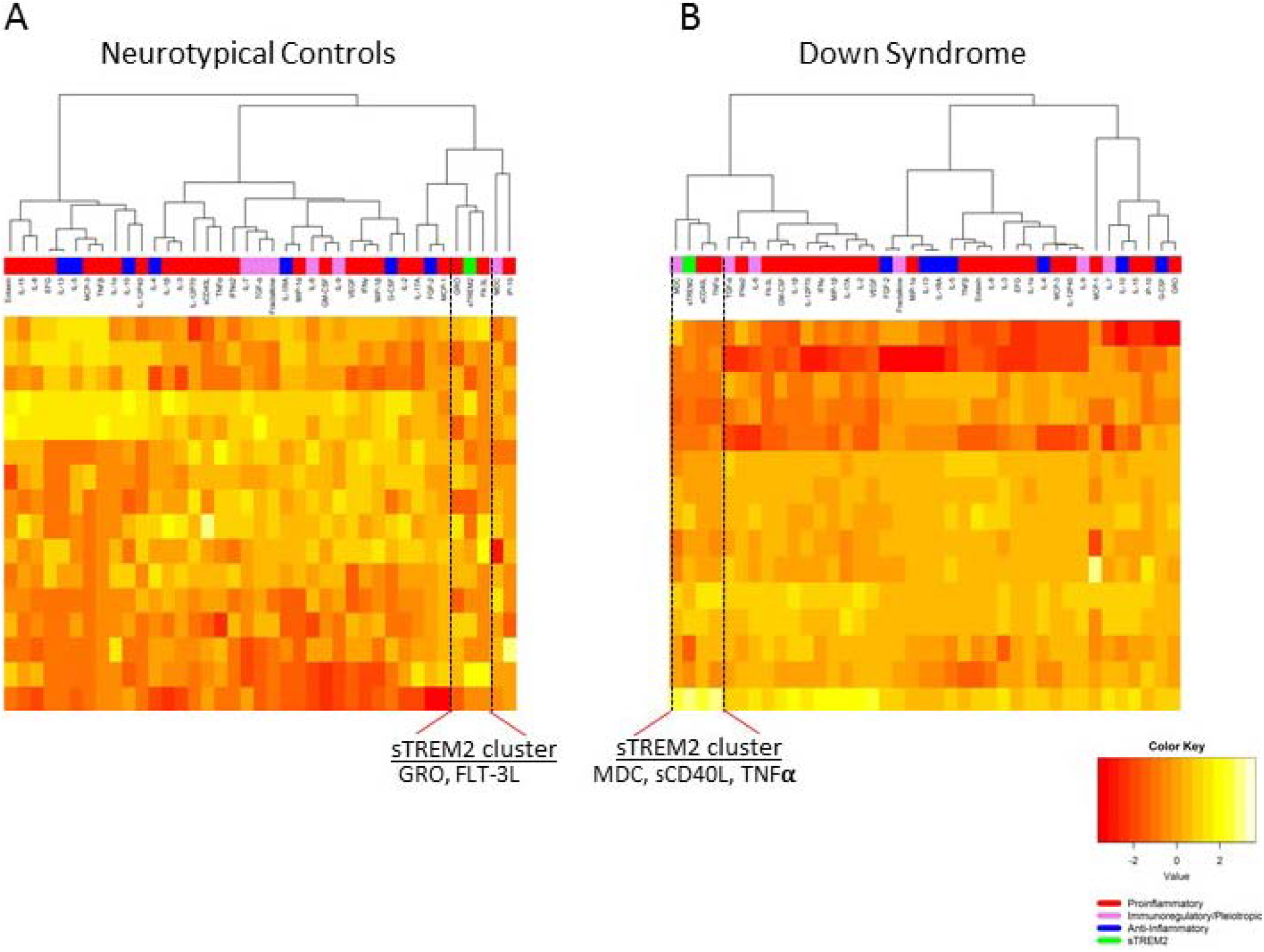
Plasma sTREM2 Clusters with MDC, sCD40L and TNFα in adults with DS, not controls. Hierarchical clustering of sTREM2 and inflammatory markers in the two groups. sTREM2 clustered with Flt-3L and GRO in the control group **(A)**, and with MDC, sCD40L, and TNFa in the group with DS **(B)**. Participants in each group, shown in the rows, were arranged based on hierarchical clustering. Inflammatory markers are separated by known functions into the general categories of anti-inflammatory (blue), immunoregulatory/pleiotropic (lavender), and pro-inflammatory (red). Individual data points that had calculated concentrations below background levels for the assay were considered physiological Os and set equal to 0.001 prior to log transform at ion for analysis purposes. Heat map values shown were scaled, log-transformed cytokine concentrations. Yellow indicates higher levels compared to red.

## 2. Discussion

In this exploratory study of young adults with DS, we found evidence of peripheral inflammation and elevated sTREM2 in a group of young adults with DS, despite the absence of AD symptoms. These participants with DS had increased levels of anti-inflammatory, proinflammatory and immunoregulatory cytokines compared to neurotypical controls, which may be explained by the body’s attempt for balance during an inflammatory state. These results cannot be explained by known acute or autoimmune illnesses in our cohort, and in fact the DS participants had lower levels of lymphocytes compared to controls. While the observed basophilia has been reported in DS ^50^, significantly decreased lymphocyte count and elevated (or trending towards elevated) neutrophils has been reported in AD ^51,52^. Our previous work showed a relationship between plasma sTREM2 and CRP in AD-related mild cognitive impairment (pre-dementia), ^27^ that was not replicated in this DS pre-dementia cohort. There might be multiple reasons for this that warrant further study. For example, since TREM2 activity has been previously described to depend on disease stage, ^53^ it is possible that this finding is related to differing stages of AD-related progression in DS pre-dementia compared to AD-related mild cognitive impairment.

Plasma sTREM2 was significantly elevated in DS, in agreement with previously described elevation of sTREM2 in DS sera from young adults with DS prior to dementia onset ^33^. Interestingly, sTREM2 is elevated in cerebrospinal fluid in the mild cognitive impairment stage of AD and in early onset forms of AD, but not in peripheral biofluids, such as plasma ^24,25,27,28,54^. The elevation of sTREM2 in young DS pre-dementia in plasma and sera suggests that peripheral sTREM2 may increase very early in individuals with elevated Aβ. Studies of AD-related biomarkers, TREM2, and other inflammatory markers would be useful in determining the potential biologic interaction between the peripheral immune system and early AD pathology. Longitudinal studies of peripheral and cerebrospinal sTREM2 in adults with DS would also further elucidate the possible role of the innate immunity in both variable onset of AD pathology and clinical dementia in DS.

We observed elevated cytokines in the group with DS compared to the neurotypical group. Our findings were similar to another study that found elevated TNFα, IL-6, and IL-10 in adults with DS pre-dementia^15^. Interestingly, in this previous report when these inflammatory markers were combined with measures of Aβ, they predicted AD development ^15^. In further support of our findings, another report describes increased IL-6, VEGF-A, MCP-1, IL-22, and TNFα, but not other cytokines, in adults aged 20-65 with DS ^55^. In contrast, a previous study of adults (mean age 30 years in DS and controls) found elevated serum MIP-1α, but not IFNγ, TNFα, MIP-1β, RANTES, or IL-6 ^56^. The similarities across these studies and our own study support the notion that in general the immune response is altered in DS, regardless of age, which may contribute to the development of AD.

In the group with DS, sTREM2 had significant positive correlations with 24 out of the 38 measured cytokines, while in the control group sTREM2 did not significantly correlate with any cytokines. Together, it was clear that people with DS had an altered relationship between sTREM2 and inflammatory markers, which could be contributing to the accelerated onset of AD in this population.

Our novel finding was that sTREM2 differed in its clustering and correlation patterns between neurotypical and DS groups. Hierarchical clustering linked sTREM2 to different inflammatory markers within the control and DS groups. Because the controls had lower levels of sTREM2, a logical conclusion is that they had more functional, membrane-bound TREM2, an important immune factor for normal innate immunity function ^19,20^. This remains to be determined, but may have important implications for the cluster analysis results observed in the control group where sTREM2 clustered with Flt-3L and GRO (**Figure 4A**). In contrast, in the DS group, sTREM2 was elevated, and clustered with inflammatory cytokines TNFα and sCD40L, and immunoregulatory cytokine MDC (**Figure 4B**). These data point to an altered function of sTREM2 when elevated, as observed in DS pre-dementia.

The clustering of sTREM2 with Flt-3L and GRO in neurotypical controls is novel information and interesting because Fms-like tyrosine kinase 3 ligand (Flt-3L), the only known ligand for Flt (CD135), is important for proliferation of hematopoietic stem cells. Knockout mice models have shown that it is crucial for the development of hematopoietic progenitor cells, B cells, and dendritic cells ^57,58^. Flt-3L was elevated in our DS pre-dementia cohort, suggesting that normally the role of TREM2 cleavage and consequent production of sTREM2 may be related to Flt-3L activity. In contrast, growth-related oncogene (GRO, also called CXCL1), although grouped with sTREM2 in controls, was not significantly elevated in our DS cohort suggesting a weaker role in DS than Flt-3L. Given that GRO is expressed in monocytes and neutrophils and has proinflammatory and mitogenic functions ^59,60^, it may have an important TREM2-related role in normal innate immunity function. TREM2 is similarly involved in cell proliferation, specifically of microglia and macrophages through an interaction with the adapter protein DNAX activating protein of 12 kDa (DAP12) ^61^.

In contrast, elevated sTREM2 in the DS cytokine secreted by activated platelets. It has been shown to be elevated in early AD and group clustered with TNFα and sCD40L, as well as with MDC. While MDC had a similar concentration in both the DS and control groups, TNFα and sCD40L were significantly elevated in DS. TNFα is an acute-phase cytokine secreted by cells including activated macrophages and brain microglial, which also express TREM2 ^57,62,63^. TNFα has been implicated in AD pathology numerous times and is reportedly associated with AD progression ^64,65^. Soluble CD40 ligand (sCD40L, or CD154) is an inflammatory there is evidence that it contributes to disease progression ^66–68^. Macrophage-derived chemokine (MDC, or CCL22) is expressed by activated T cells, NK cells, macrophages, and monocytes, and has chemoattractant and inflammatory properties ^69^. Few studies to date have looked at MDC in the context of AD, but one study found that levels of MDC in CSF decreased after one year of resveratrol treatment in AD compared to the placebo group ^70^, which may implicate MDC in neuroinflammatory pathways. Given that our cohort of young adults with DS were all pre-dementia, these observed correlations with sTREM2 may be early predictors of dementia onset.

This is an exploratory study that is limited by the small sample size and potential for false negative results. DS biomarker studies, including our study, are limited by samples size and should be considered with caution until replicated in larger populations ^10,15,71,72^. Covariates such as age and *APOE*4 status did not differ between our two groups but are important to control for in larger future studies. Another limitation of this study is that karyotypes were not available for the DS participants. Although unlikely, given the unique clinical presentation of these two types of patients, it is possible that some of the DS participants had mosaic DS or chromosome 21 translocations, rather than full trisomy 21. This could lead to heterogeneity of the subjects. Additionally, we were limited by the availability of samples from this group. We were not able to collect CSF from these subjects. CSF has been studied more in-depth in AD studies compared to plasma, and could lend additional evidence of the contribution of sTREM2 and the observed immune state in DS. Furthermore, we were not able to perform any functional studies with peripheral cells in our cohort. Future studies of *ex vivo* peripheral blood cell production of sTREM2 and the relationship with the identified inflammatory markers will allow us to understand whether sTREM2 is driving the inflammatory state, or counteracting it. For these reasons, a larger replication study would be important to help support results from this exploratory study.

## 3. Conclusion

The results of this exploratory study strongly suggest a relationship between plasma sTREM2 and inflammatory markers in DS pre-dementia. We observed significantly elevated inflammatory markers in DS, in agreement with previous reports ^16,18,39,73^. Notably, we observed a strong positive correlation between many of the tested immune factors and sTREM2 in the group with DS that was not observed in controls. To our knowledge, this is the first report of a correlation between sTREM2 and multiple inflammatory markers in DS. This novel result is supported by previously described dysregulation of the immune response in DS ^16,18,39,73^ and suggests that TREM2 plays a role in the peripheral immune response in DS pre-dementia.

## Supplemental Figures

**Supplemental Table 1.**
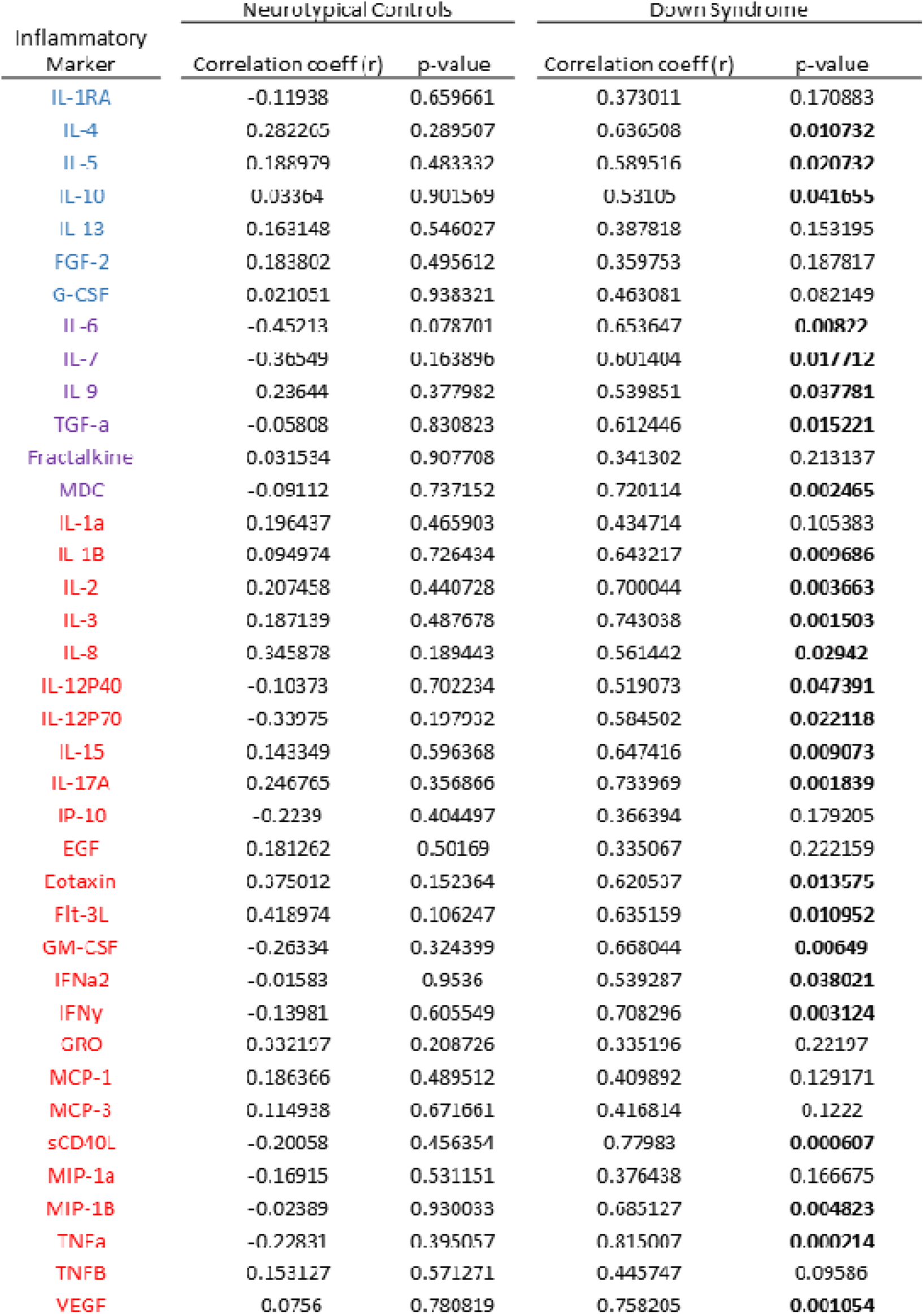
The relationship between peripheral sTREM2 and peripheral inflammatory markers in adults with Down Syndrome pre-dementia compared to neurotypical controls. There were significant correlations (Pearson correlation not corrected for multiple comparisons) between plasma sTREM2 and plasma inflammatory markers in each group. P-values <0.05 are bolded.

**Supplemental Figure 1.**
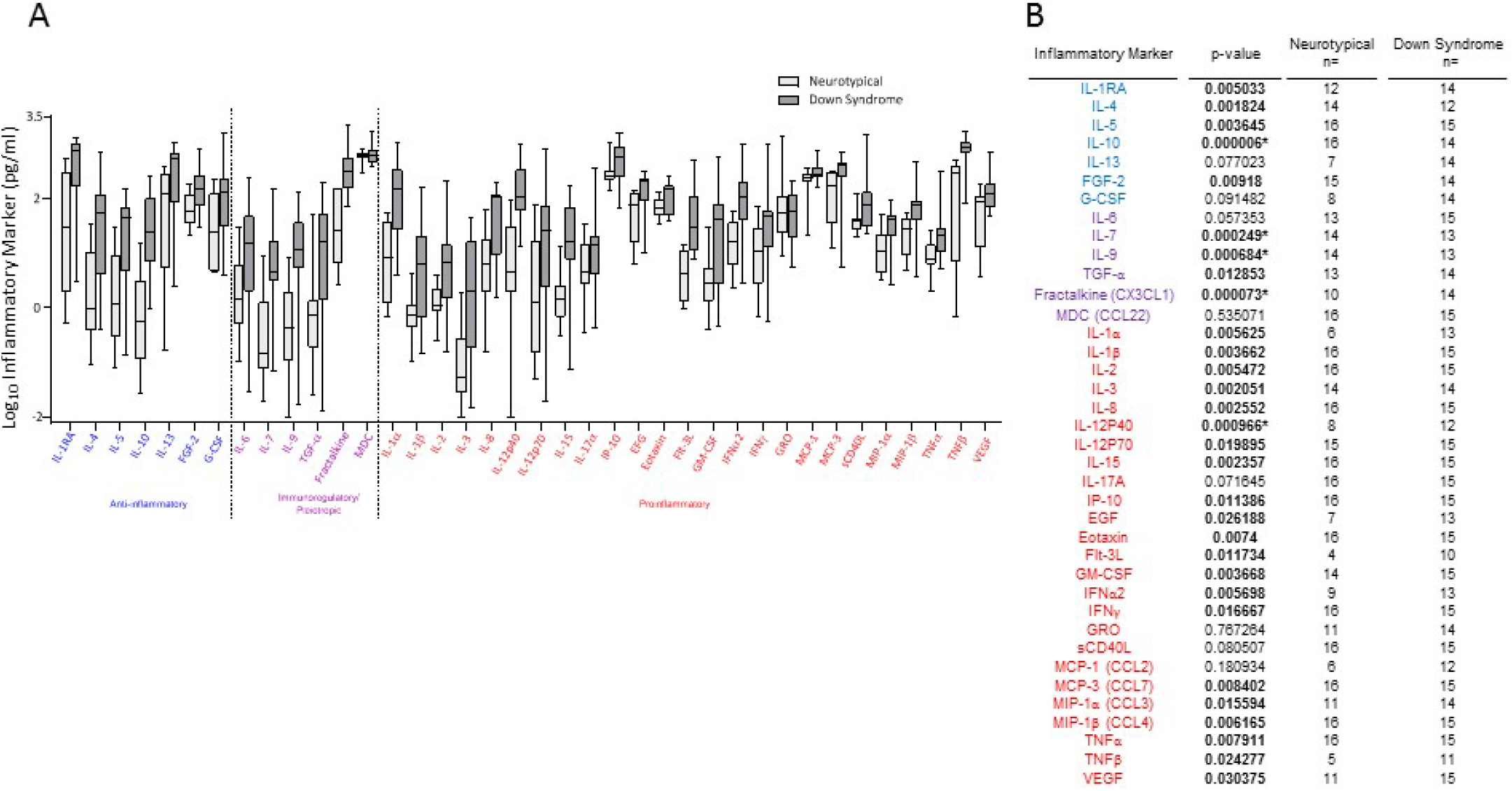
Plasma Inflammatory Markers in Down Syndrome Pre-dementia (removing values below background detection). Significantly higher plasma inflammatory markers in young adults with Down Syndrome pre-dementia (dark grey bars), compared to neurotypical controls (light grey bars) for 30/38 of the inflammatory markers on the panel **(A)**, significance as indicated by unpaired t-test p-values (asterisk denotes 1% FDR significance) **(B)**. Inflammatory markers are separated by known functions into the general categories of anti-inflammatory (blue), immunoregulatory/pleiotropic (purple), and pro-inflammatory (red). Individual data points that had calculated concentrations below background levels for the assay were considered physiological Os and removed from the analysis. Participants analyzed per group differed based on cytokine due to the number of values below background, as noted in B. Lines indicate mean levels. Minimum and maximum levels are indicated.

## Abbreviations

AD: Alzheimer’s disease
DS: Down syndrome
TREM2: Triggering receptor expressed in myeloid cells 2
Aβ: amyloid-β
CBC: Complete blood counts
ESR: erythrocyte sedimentation rate

